# Frameshift and wild-type proteins are highly similar because the genetic code and genomes were optimized for frameshift tolerance

**DOI:** 10.1101/067736

**Authors:** Xiaolong Wang, Quanjiang Dong, Gang Chen, Jianye Zhang, Yongqiang Liu, Yujia Cai

## Abstract

Frameshift protein sequences encoded by alternative reading frames of coding genes have been considered meaningless, and frameshift mutations have been considered of little importance for the molecular evolution of coding genes and proteins. However, functional frameshifts have been found widely existing. It was puzzling how a frameshift protein kept its structure and functionality while its amino-acid sequence was changed substantially. Here we show that frame similarities between frameshifts and wild types are higher than random similarities and are defined at the genetic code, gene, and genome levels. In the standard genetic code, frameshift codon substitutions are more conservative than random substitutions. The frameshift tolerability of the standard genetic code ranks in the top 2.0-3.5% of alternative genetic codes, showing that the genetic code is nearly optimal for frameshift tolerance. Furthermore, frameshift-resistant codons (codon pairs) appear more frequently than expected in many genes and certain genomes, showing that the frameshift optimality is reflected not only in the genetic code but more importantly, in its allowance of further optimizing the frameshift tolerance of a particular gene or genome, which shed light on the role of frameshift mutations in molecular and genomic evolution.

## 1. Background

The genetic code was deciphered in the 1960s [1]. The standard genetic code consists of 64 triplet codons, 61 sense codons for the twenty amino acids (AAs), and three nonsense codons for stop signals. The natural genetic code has several important properties: (1) the genetic code is universal in all species, with only a few variations found in some organelles or organisms, such as mitochondrion, archaea, yeast, and ciliates [2]; (2) the triplet codons are redundant, degenerate, and the third base is wobble (interchangeable); (3) in a coding DNA sequence (CDS), an insertion or deletion (InDel) causes a frameshift mutation if its size is not a multiple of three.

It has been reported that the natural genetic code was optimized for translational error minimization, which is being extremely efficient at minimizing the effect of point mutation or mistranslation errors and is optimal for kinetic energy conservation in polypeptide chains [3-6]. Moreover, it was discovered that the standard genetic code resists frameshift errors by increasing the probability that a stop signal is encountered upon frameshifting because frameshifted codons for abundant amino acids overlap with stop codons [7].

A frameshift mutation alters a reading frame, producing frameshift protein sequences (*frameshifts*). Frameshifts have long been considered mostly meaningless since they look completely different from the wild type and are often interrupted by many stop signals. A frameshifted gene yields truncated, non-functional, and potentially cytotoxic peptides [8]. Therefore, frameshift mutations have been considered harmful and of little importance to the evolution of proteins or coding genes. However, it is known that frameshifting does not always lead to lost-of-function. Frameshifted genes can sometimes be expressed through special mechanisms, such as translational readthrough [9-11], ribosomal frameshifting [12-14], reading frame transition [13], or genetic recoding [15]. Moreover, frameshifted genes can be retained for millions of years and enable the acquisition of new functions [16].

Many cases of functional frameshift homologs have been reported [17-19], *e*.*g*., by collecting human coding exons bearing InDels compared with the chimpanzee genome, *Hahn* and *Lee* identified nine frameshift homologs between humans and chimpanzee, some of which seem to be functional in both species [19]. It has also been reported that several functional frameshifts as compared to their related genes in other species [20]. Particularly, Bartonek et al [21] showed that frameshifting preserves key physicochemical properties of proteins; Huang et al [22] showed that frameshifted proteins of a bacteria toxin gene retain the same function. Moreover, it has been reported that frameshifting may lead to functional divergence [16], novel genes [17], or overlapping genes, in viruses [23], bacteria [24], and even humans [25].

As is well known, a protein can be dysfunctioned even by changing one residue, so it is very puzzling how a frameshift protein kept its tertiary structural and functional integrity while its primary sequence was changed substantially. We have consistently observed high similarities among frameshifts and wild-type protein sequences [26], while our previous analyses based on ClustalW alignments were defective. Since we disclosed this study, other groups have further analyzed the genetic code using the physicochemical properties (PCPs) of the amino acids [27]. Here, we reanalyze the data using a novel frameshift alignment method and report that frameshifts and wild types are always highly similar and that the genetic code is nearly optimal for frameshift tolerance. Furthermore, many genes and certain genomes were further optimized to enhance their tolerance to frameshift mutations, which shed light on the role of frameshift mutations in molecular and genomic evolution.

A frameshift mutation alters the reading frame of a gene and produces frameshifted proteins (*frameshifts*). Frameshifts have long been considered meaningless because they look completely different from the wild type. However, many cases of functional frameshifts have been widely observed. It was puzzling how a frameshift protein maintains its structure and functionality. Here we show that the similarities between frameshifts and their wild types are significantly higher than expected. We demonstrate that the genetic code is nearly optimal in terms of frameshift tolerance, making it prevail in early evolution. More importantly, it allows further optimizing of a particular gene or genome to tolerate frameshift mutations and sheds light on the role of frameshift mutations in molecular and genomic evolution.

## 2. Materials and Methods

### 2.1 Protein-coding DNA sequences

All reference coding sequences (CDSs) in ten model species, including *Escherichia coli, Saccharomyces cerevisiae, Arabidopsis thaliana, Caenorhabditis elegans, Drosophila melanogaster, Danio rerio, Xenopus tropicalis, Mus musculus, Pan troglodytes*, and *Homo sapiens*, were retrieved from UCSC, Ensembl, or NCBI Genome Databases. Ten thousand sets of CDSs, each containing three CDSs with 300 or 500 random sense codons, were produced by a homemade program (RandomCDSs.java).

### 2.2 Aligning and computing the similarities of wild types and frameshifts

Program Similarity.java batch translates CDSs and computes the pairwise similarities among the translations, in which CDSs are translated using the standard genetic code in the 3 different reading frames in the sense strand, and the 3 different translations are aligned by 3 different methods, including *ClustalW2, MSA*, or *FrameAlign*. To calculate pairwise similarity, a pair of matched AAs in a pairwise alignment is considered conserved if their substitution score is ≥ 0 in the scoring matrix GON250, *i.e*., gaps and negative scores are considered different. The percent of conserved sites gives the pairwise similarity between a frameshift and the corresponding wild-type protein sequence.

Similarity.java translates internal stop codon into AAs using a set of readthrough rules (Table 1). Translational readthrough occurs upon the suppressor tRNA activity with an anticodon matching a stop codon [11]. Many studies showed that translational readthrough occurs in prokaryotes and eukaryotes, from *E. coli* to humans, while the readthrough rules may vary among different species [28]. In *E. coli*, nonsense suppression tRNAs reported includes amber suppressors (*supD* [29], *supE* [30], *supF* [31]), ochre suppressors (*supG* [32]), and opal suppressors (*supU* [31], *su9* [33]). In this study, suppressor tRNAs were summarized as a set of readthrough rules and used to translate frameshifted CDSs.

**Table 1.**
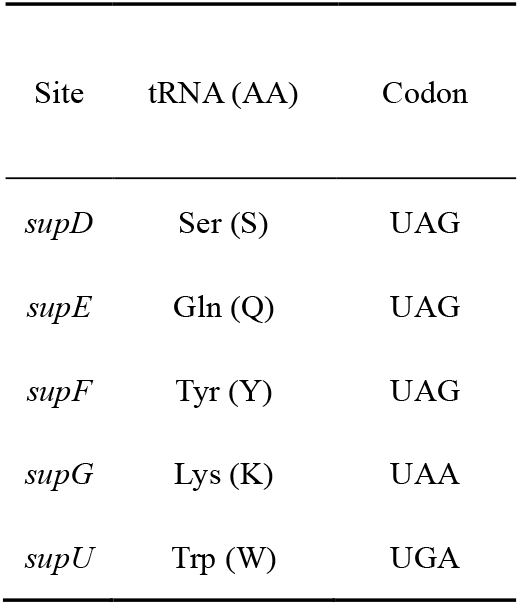
The *readthrough rules* derived from natural suppressor tRNAs for nonsense mutations.

| Site | tRNA (AA) | Codon |
| --- | --- | --- |
| *supD* | Ser (S) | UAG |
| *supE* | Gln (Q) | UAG |
| *supF* | Tyr (Y) | UAG |
| *supG* | Lys (K) | UAA |
| *supU* | Trp (W) | UGA |

### 2.3 FrameAlign: aligning of frameshifts and wild-type protein sequence

A wild-type protein-coding gene sequence consists of *n* triplet codons is written as:

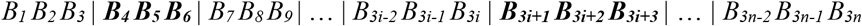

Where *B*_*k*_ ∈{*A, G, U, C*}; *i* = 1… *n*; *k =* 1…3*n*. Each pair of neighboring codons are separated by a bar to show the reading frame. Its encoded wild-type protein sequence (*WT*), consisting of *n* amino acids, can be written as,

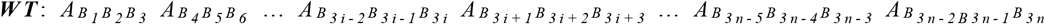

where 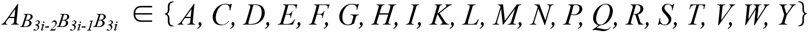,represents the amino acid encoded by the *i*^*th*^ codon (*B*_*3i-2*_ *B*_*3i-1*_ *B*_*3i*_). If a frameshift is caused by deleting or inserting one or two bases in the start codon, there are only four cases:

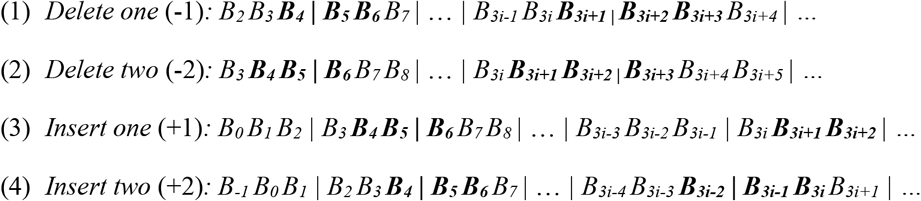

If a frameshift mutation occurs at any location between the first and the *i*^*th*^ codon, the (*i*+1)^*th*^ codon (***B***_***3i+1***_ ***B***_***3i+2***_ ***B***_***3i+3***_) has only two possible changes:

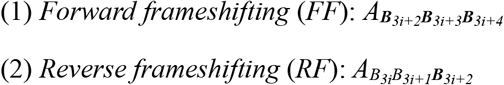

This continues for each codon downstream, resulting in two frameshifts, denoted as *FF* and *RF*,

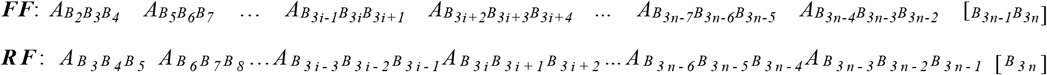

The final codon of *FF* or *RF*, as shown in the square brackets, is incomplete and was deleted. The *i*^*th*^ codon of the frameshifts, *B*_*3i+2*_ *B*_*3i+3*_*B*_*3i+4*_ (*FF*) and *B*_*3i*_ *B*_*3i+1*_*B*_*3i+2*_ (*RF*), both have two bases overlapping with the (*i*+1)^*th*^ *WT* codon, *B*_*3i+1*_ *B*_*3i+2*_ *B*_*3i+3*_, and their encoded amino acids, 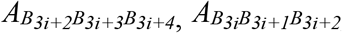 and 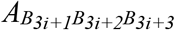 are likely similar to each other because similar codons encode amino acids with related physicochemical properties [3]. As shown in the following, *WT, FF*, and *RF* can be aligned in pairs, called *FrameAlign*, but cannot be aligned properly in a multiple sequence alignment (MSA), so common MSA tools are not suitable for aligning wild-type and frameshifts.

1. ***WT vs. FF:*** insert one gap at the end of *FF*.

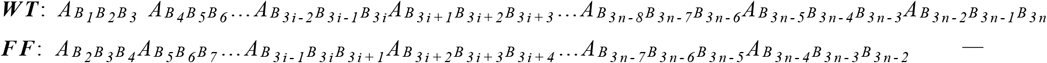
2. ***WT vs. RF:*** insert one gap at the beginning of *RF*.

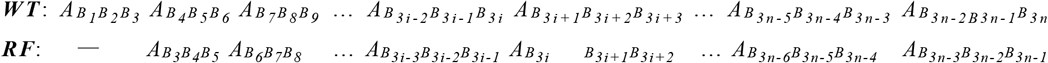
3. ***FF vs. RF*:** no gaps are needed.

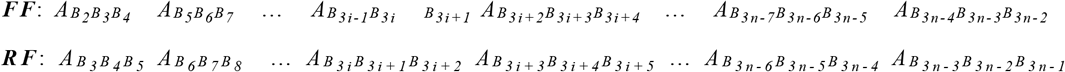

### 2.4 Computational analysis of frameshift codon substitutions

According to whether the encoded AA is changed or not, codon substitutions have been classified into *synonymous substitutions* (SSs) and *nonsynonymous substitutions* (NSSs). Based on the above analysis in section 2.3, we further classified codon substitutions into three subtypes:

1. *Random substitutions (RCSs)*: randomly change all three bases of the codons, including 64 × 64 = 4096 possible codon substitutions.
2. *Wobble substitution (WCSs)*: randomly change only the third position of the codons, including 64 × 4 = 256 possible codon substitutions.
3. *Frameshift substitution* (*FCSs*): codon substitutions caused by forward or reverse frameshifting. Each codon has 4 forward and 4 reverse FCSs, and there are 64 × 8 = 512 FCSs in total.

In most cases, all three bases in the frameshifted codon are changed compared with the original codon, except for triplet monomers (such as AAA, GGG). The AA substitution scores of FCSs and RCSs are defined as frameshift substitution scores (FSSs) and random substitution scores (RSSs), respectively. The sum FSS for all possible FCSs is considered the frameshift tolerability of the genetic code. Program Frameshift-CODON.java computes the substitution score for each codon substitution by using a scoring matrix, BLOSSUM62 [34], PAM250 [35, 36], or GON250 [37].

### 2.5 Computational analysis of random or alternative codon tables

RandomCodes.java generates random codon tables by swapping AAs assigned to the sense codons and keeping all degenerative codons synonymous (*Freeland* and *Hurst* [5]). One million random codon tables were sampled from all possible (20! = 2.43290201×10^18^) genetic codes randomly using a random-number-based sampling algorithm, in which the probability of an AA being swapped is proportional to its proportion in the code table. The sampling was repeated 100 times independently. For each sample, the sum of FSSs for each genetic code was computed and compared with that of the natural genetic code.

AlternativeCodes.java produces all (13824) alternative codon tables by permuting the nucleotide in each codon position independently (*Itzkovitz* and *Alon* [7]). Each alternative code has the same number of codons per amino acid and the same impact of misread errors as in the standard genetic code. The sum of FSSs for each of the compatible genetic codes was computed and compared with that of the natural genetic code.

### 2.6 Analysis of codon pairs and their frameshift substitution scores

FrameshiftCodonPair.java computes the FSSs for all possible codon pairs. For a given codon pair, written as *B*_*1*_ *B*_*2*_ *B*_*3*_ | ***B***_***4***_ ***B***_***5***_ ***B***_***6***_, its encoded AA pair is written as 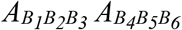. There are 400 different AA pairs, 64 × 64 = 4096 different codon pairs. Similarly, the codon pair and its encoded AAs have only two types of changes in frameshifting:

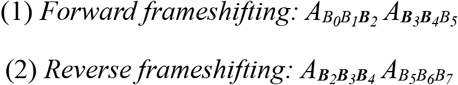

where *B*_*0*_ and *B*_*7*_ each have four choices, and therefore, there are 4096 × 8 = 32,768 different frameshift codon pair substitutions (FCPSs). For each FCPSs, 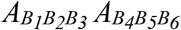 was compared with their frameshifts to obtain their FSSs.

### 2.7 Computational analysis of the usage of codon and codon pairs

For each genome, the number of occurrences was counted for every codon or codon pair. The observed and expected frequencies were then calculated for each codon or codon pair using *Gutman* and *Hatfield* method [38]. These calculations result in a list of 64 codons and 4096 codon pairs, each with an expected (*E*) and observed (*O*) number of occurrences, frequency, together with a value for the *χ*^2^ statistics. A codon or codon pair was identified as over-represented if *O* > *E* (or under-represented if *O* < *E*), and the average FSSs were calculated for each genome weighted by their codon or codon pair usages.

## 3. Results and Analysis

### 3.1 Wild-type and frameshift protein sequences are always highly similar

First, 100,000 random CDSs each containing 300 sense codons were simulated and translated into protein sequences in the three reading frames in the sense strand. The three translations were aligned using *ClustalW, MSA*, or *FrameAlign*, and their *frame similarities* and *random similarities* were calculated, respectively. Similarities among the translations of three different reading frames are defined as frame similarities and those among the translations of three different random CDSs as random similarities. Frame similarities were also calculated for all available real CDSs in ten model organisms.

When translations were aligned using ClustalW, the estimated average frame similarity for real and random CDSs is 0.456±0.033 and 0.452±0.013 (Table 2a), respectively. However, on average, ClustalW placed 49.57 and 80.11 gaps in the alignments of the translations of real and random CDSs, respectively. Besides, the estimated average random similarity is comparable to the average frame similarity but on average 137.05 gaps were placed in the alignments of translations of random CDSs, indicating that the similarity calculations can be false due to alignment artifacts, caused by inserting excessive gaps.

**Table 2a.**
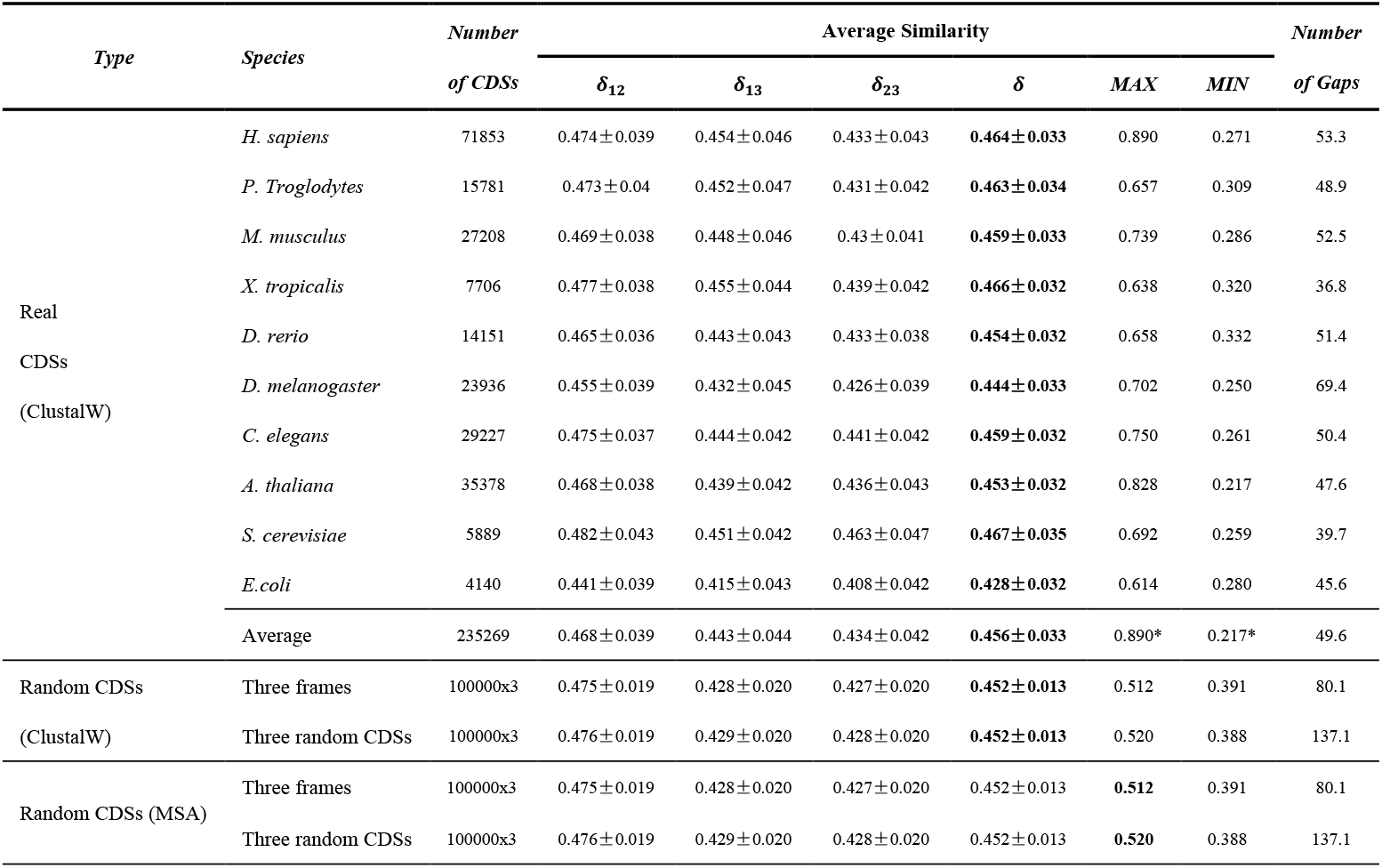
The similarities of proteins and their frameshifts (aligned by *ClustalW* or *MSA*).

| Type | Species | Number<br>of CDSs | Average Similarity |  |  |  |  |  | Number<br>of Gaps |
| --- | --- | --- | --- | --- | --- | --- | --- | --- | --- |
| | | | $\delta_{12}$ | $\delta_{13}$ | $\delta_{23}$ | $\delta$ | MAX | MIN | |
| Real<br>CDSs<br>(ClustalW) | *H. sapiens* | 71853 | 0.474±0.039 | 0.454±0.046 | 0.433±0.043 | **0.464±0.033** | 0.890 | 0.271 | 53.3 |
|  | *P. Troglodytes* | 15781 | 0.473±0.04 | 0.452±0.047 | 0.431±0.042 | **0.463±0.034** | 0.657 | 0.309 | 48.9 |
|  | *M. musculus* | 27208 | 0.469±0.038 | 0.448±0.046 | 0.43±0.041 | **0.459±0.033** | 0.739 | 0.286 | 52.5 |
|  | *X. tropicalis* | 7706 | 0.477±0.038 | 0.455±0.044 | 0.439±0.042 | **0.466±0.032** | 0.638 | 0.320 | 36.8 |
|  | *D. rerio* | 14151 | 0.465±0.036 | 0.443±0.043 | 0.433±0.038 | **0.454±0.032** | 0.658 | 0.332 | 51.4 |
|  | *D. melanogaster* | 23936 | 0.455±0.039 | 0.432±0.045 | 0.426±0.039 | **0.444±0.033** | 0.702 | 0.250 | 69.4 |
|  | *C. elegans* | 29227 | 0.475±0.037 | 0.444±0.042 | 0.441±0.042 | **0.459±0.032** | 0.750 | 0.261 | 50.4 |
|  | *A. thaliana* | 35378 | 0.468±0.038 | 0.439±0.042 | 0.436±0.043 | **0.453±0.032** | 0.828 | 0.217 | 47.6 |
|  | *S. cerevisiae* | 5889 | 0.482±0.043 | 0.451±0.042 | 0.463±0.047 | **0.467±0.035** | 0.692 | 0.259 | 39.7 |
|  | *E.coli* | 4140 | 0.441±0.039 | 0.415±0.043 | 0.408±0.042 | **0.428±0.032** | 0.614 | 0.280 | 45.6 |
|  | Average | 235269 | 0.468±0.039 | 0.443±0.044 | 0.434±0.042 | **0.456±0.033** | 0.890* | 0.217* | 49.6 |
| Random CDSs<br>(ClustalW) | Three frames | 100000x3 | 0.475±0.019 | 0.428±0.020 | 0.427±0.020 | **0.452±0.013** | 0.512 | 0.391 | 80.1 |
|  | Three random CDSs | 100000x3 | 0.476±0.019 | 0.429±0.020 | 0.428±0.020 | **0.452±0.013** | 0.520 | 0.388 | 137.1 |
| Random CDSs (MSA) | Three frames | 100000x3 | 0.475±0.019 | 0.428±0.020 | 0.427±0.020 | 0.452±0.013 | **0.512** | 0.391 | 80.1 |
|  | Three random CDSs | 100000x3 | 0.476±0.019 | 0.429±0.020 | 0.428±0.020 | 0.452±0.013 | **0.520** | 0.388 | 137.1 |

To sidestep the effect of aligners, *MSA* was used to obtain the optimal alignments [39]. Unfortunately, because of the memory requirements, *MSA* cannot be applied to protein sequences >500 AAs, so it cannot be applied to many real genes. So, only the translations of random CDSs were aligned using *MSA*, and the estimated average frame similarity is 0.410±0.055 (Table 2a); on average, *MSA* placed 108.3 gaps in the alignments. However, the estimated average random similarity is also as high as 0.412±0.055, and on average, *MSA* placed 109.5 gaps in the alignments of random protein sequences, suggesting that false similarities caused by gappy alignment artifacts cannot be avoided using optimal alignments.

As described in section 2.3, owing to inherent constraints, frameshifts and wild types cannot be aligned correctly using any existing methods. We designed *FrameAlign*, a simple method for pairwise alignment of frameshifts and wild types. For example, in a FrameAlign of wild-type zebrafish VEGFAA and its frameshifts, the average amino-acid sequence similarity is as high as 52.34% (Fig 1). This is very surprising, so we must emphasize here that this case was not cherry-picked but arbitrarily selected. One could reproduce the same kind of results easily with almost any real coding genes.

**Fig 1.**
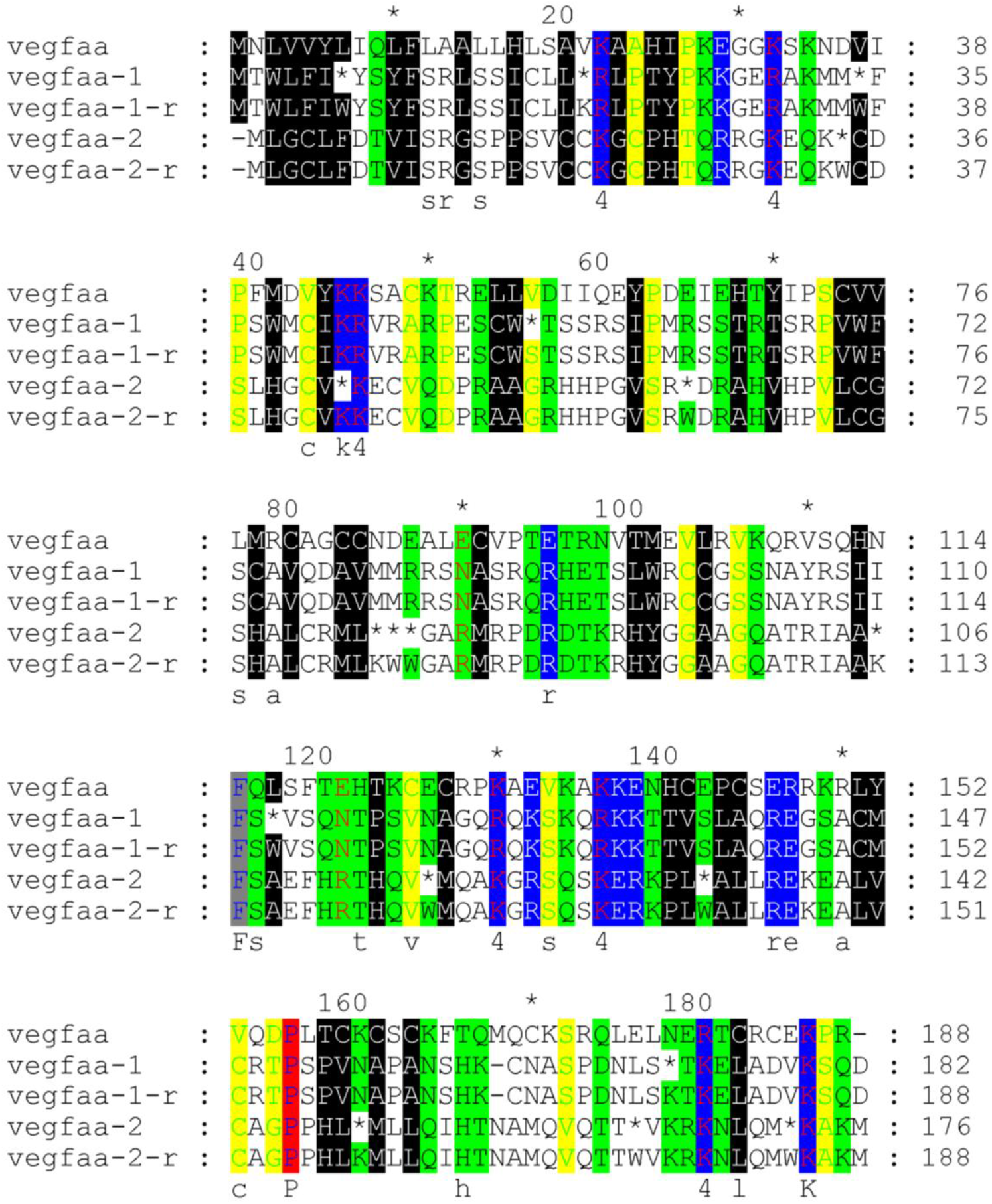
The alignment of wild-type VEGFAA, readthrough, or non-readthrough translation of the frameshifts. Vegfaa: wild-type VEGFAA; vegfaa-1: -1 frameshift non-readthrough translation; vegfaa-2: -2 frameshift non-readthrough translation; vegfaa-1-r: -1 frameshift readthrough translation; vegfaa-2-r: -2 frameshift readthrough translation;

**Fig 2.**
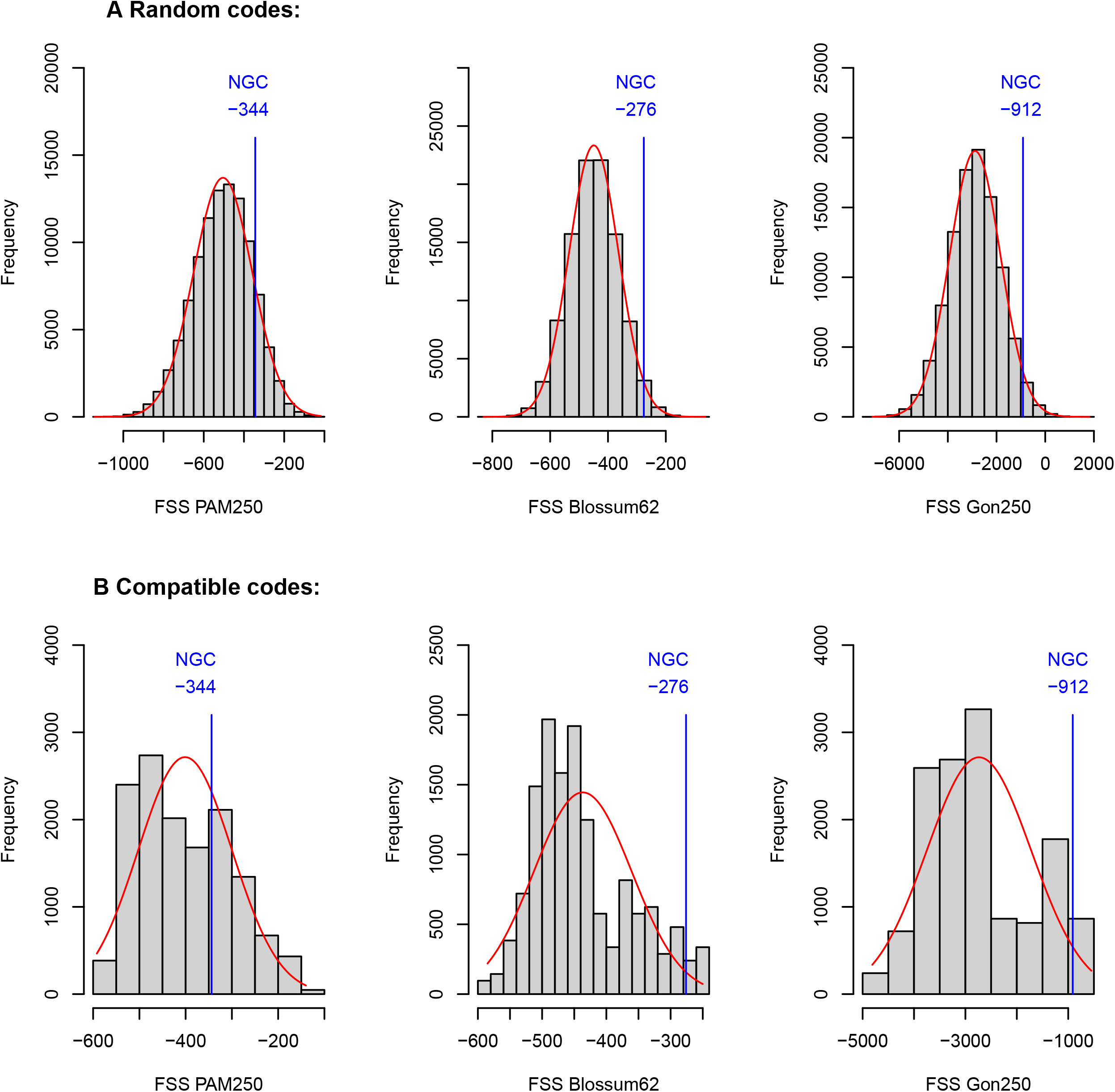
The distribution of the FSSs for the alternative genetic codes. A) randomly chosen one million random codon tables and (B) all 13824 alternative codon tables. NGC: the natural genetic code; FSSs were calculated using matrices PAM250, BLOSSUM62, and GON250. The probability densities were computed using a normal distribution function and plotted in language R.

When the translations were aligned using *FrameAlign*, the estimated average random similarity is 0.383±0.018, and the mean frame similarity is 0.394±0.016 (Table 2b). Their difference is small but statistically extremely significant (t-test P-value ≈ 0). Furthermore, the overall mean frame similarity of the real genes is as high as 0.450±0.030 (Table 2b, S1), much higher than random similarity (t-test P-value ≈ 0), or the frame similarity of random CDSs (t-test P-value ≈ 0), indicating that frameshifts of real genes are even more like their wild types, which cannot be revealed by calculations based on *ClustalW* or *MSA* alignments.

For a given CDS, let *δ*_*ij*_ be the pairwise similarities of its three translations, *i, j=1,2,3, i ≠ j, δ*_*ij*_ = *δ*_*ji*_. Using *FrameAlign*, the average similarity among the frameshifts and the wild type is defined as *the shiftability of protein-coding genes* (*δ*),

**Table 2b.** The similarities of proteins and their frameshifts (aligned by *FrameAlign*)

| Type | Species | Number<br>of CDSs | Average Similarity |  |  |  |  |  | Number<br>of Gaps |
| --- | --- | --- | --- | --- | --- | --- | --- | --- | --- |
| | | | $\delta_{12}$ | $\delta_{13}$ | $\delta_{23}$ | $\delta$ | MAX | MIN | |
| Real<br>CDSs<br>(FrameAlign) | *H. sapiens* | 71853 | 0.492±0.043 | 0.472±0.044 | 0.434±0.040 | **0.466±0.029** | 0.713 | 0.194 | 2 |
|  | *P. Troglodytes* | 15781 | 0.491±0.046 | 0.468±0.046 | 0.431±0.042 | **0.463±0.030** | 0.625 | 0.311 | 2 |
|  | *M. musculus* | 27208 | 0.484±0.046 | 0.469±0.042 | 0.426±0.040 | **0.460±0.029** | 0.739 | 0.286 | 2 |
|  | *X. tropicalis* | 7706 | 0.481±0.042 | 0.481±0.041 | 0.439±0.037 | **0.467±0.028** | 0.644 | 0.353 | 2 |
|  | *D. rerio* | 14151 | 0.471±0.044 | 0.468±0.040 | 0.408±0.040 | **0.449±0.030** | 0.614 | 0.314 | 2 |
|  | *D. melanogaster* | 23936 | 0.475±0.046 | 0.457±0.044 | 0.362±0.047 | **0.431±0.030** | 0.689 | 0.236 | 2 |
|  | *C. elegans* | 29227 | 0.450±0.047 | 0.475±0.045 | 0.421±0.043 | **0.449±0.032** | 0.634 | 0.224 | 2 |
|  | *A. thaliana* | 35378 | 0.442±0.045 | 0.477±0.044 | 0.412±0.041 | **0.444±0.031** | 0.882 | 0.244 | 2 |
|  | *S. cerevisiae* | 5889 | 0.461±0.041 | 0.510±0.042 | 0.423±0.038 | **0.465±0.029** | 0.692 | 0.259 | 2 |
|  | *E.coli* | 4140 | 0.435±0.046 | 0.426±0.047 | 0.372±0.043 | **0.411±0.030** | 0.571 | 0.237 | 2 |
|  | Average | 235269 | 0.468±0.045 | 0.470±0.043 | 0.413±0.041 | **0.450±0.030** | 0.882* | 0.194* | 2 |
| Random CDSs<br>(FrameAlign) | Three frames | 100000 | 0.394±0.028 | 0.394±0.028 | 0.395±0.028 | **0.394±0.016** | 0.477 | 0.330 | 2 |
|  | Three random CDSs | 100000x3 | 0.383±0.028 | 0.383±0.028 | 0.383±0.028 | **0.383±0.018** | 0.458 | 0.304 | 0 |

\* Very large/small similarity values were observed in a few very short or repetitive peptides.

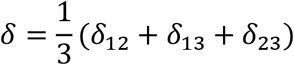

Shiftability is a quantitative measurement of frameshift tolerability. As frameshifting occurs between any two of the three reading frames, *δ*_12_, *δ*_13_, and *δ*_23_ are all considered in the formula. As shown in Table 2b, the average shiftability is close to 0.45 in real genes but less than 0.4 in random CDSs. In other words, on average, about 45% of the AA sites remain conserved in a frameshift of a real gene. As shown in Table 2b, the shiftability varies substantially in different species, from 0.411 (*E. coli*) to 0.466 (human), but the standard deviations are all as low as 0.030 in all the species tested, *i*.*e*., shiftability is species-dependent, and *δ* is centered at a specific value for most genes in a specific species.

### 3.2 The genetic code was optimized for frameshift tolerance

As described in section 2.5, the averages of AA substitution scores for random, wobble, and frameshift substitutions were computed, respectively. As shown in Table 3 and Supp S2, in all 4096 random substitutions, only a small proportion (230/4096=5.6%) of them are synonymous, and the proportion of positive substitutions (with a positive AA substitution score) is 859/4096=20.1%. Wobble substitutions have the highest average score because most (192/256=75%) wobble substitutions are synonymous, and most (192/230=83%) synonymous substitutions are wobble. In contrast, only a small percentage (28/512=5.5%) of the frameshift substitutions are synonymous (Table 4), while the remaining 94.5% are nonsynonymous. However, 29.7% of frameshift substitutions are positive nonsynonymous, which is about 1.5-fold of that of random (20.1%) and about 2-fold of that of wobble substitutions (15.6%). In summary, in the standard genetic code, wobble substitutions are assigned mostly with synonymous AAs, while frameshift substitutions are more frequently with positive nonsynonymous ones.

**Table 3.** The amino acid substitution scores for different kinds of codon substitutions.

| *Codon Substitution* |  | *Random* | *Frameshift* |  | *Wobble* |
| --- | --- | --- | --- | --- | --- |
|  |  |  | *FF* | *RF* |  |
| *Type of*<br>*Codon*<br>*Substitution* | *All* | 4096 | 256 | 256 | 256 |
|  | *Unchanged (%)* | 64 (1.6%) | 4 (1.6%) | 4 (1.6%) | 64 (25%) |
|  | *Changed (%)* | 4032 (98.4%) | 252 (98.4%) | 252 (98.4%) | 192 (75%) |
|  | *SS (%)* | 230 (5.6%) | 14 (5.5%) | 14 (5.5%) | 192 (75%) |
|  | *NSS-Positive (%)* | 859 (20.1%) | 76 (29.7%) | 76 (29.7%) | 40 (15.6%) |
|  | *NSS-Negative (%)* | 3007 (73.4%) | 166 (64.8%) | 166 (64.8%) | 24 (9.4%) |
| *Average* | *BLOSSUM62* | -1.29 | -0.61 | -0.65 | 3.77 |
| *Substitution* | *PAM250* | -4.26 | -0.84 | -0.84 | 3.68 |
| *Score* | *GON250* | -10.81 | -1.78 | -1.78 | 35.60 |

SS/NSS: synonymous/nonsynonymous substitution; FF/RF: forward/reverse frameshift substitutions.

**Table 4.** The frameshift substitution scores of the natural and alternative genetic codes.

| *Genetic codes*<br>(*Number tested*) | *Scoring*<br>*Matrix* | *FSS of the natural genetic code* |  |  |  |  | *FSS of the alternative genetic codes* |  |  |
| --- | --- | --- | --- | --- | --- | --- | --- | --- | --- |
|  |  | *FSS Score* | *Rank* | *Rank%* | *STDEV* | *STDEV%* | *Average* | *STDEV* | *STDEV%* |
| Random<br>(1,000,000 × 100) | PAM250 | -344 | 132,586.79 | 13.26% | 1,011.17 | 0.1011% | -504.88 | 0.54 | -0.1073% |
|  | Blossum62 | -276 | 19,752.52 | 1.98% | 295.17 | 0.0295% | -450.53 | 0.27 | -0.0598% |
|  | Gonnet250 | -912 | 29,447.26 | 2.94% | 398.72 | 0.0399% | -2,872.95 | 4.16 | -0.1447% |
| Compatible<br>(13824) | PAM250 | -344 | 4273 | 30.91% | - | - | -401.25 | - | - |
|  | Blossum62 | -276 | 481 | 3.48% | - | - | -436.75 | - | - |
|  | Gonnet250 | -912 | 481 | 3.48% | - | - | -2,736.13 | - | - |

Besides, no matter which AA substitution scoring matrix (BLOSSUM62, PAM250, or GON250) is used, the average FSSs are always significantly higher than those of random substitutions. Using GON250, *e*.*g*., the average FSS (−1.78) is significantly higher than the average RSS (−10.81). As shown in Table S2, AAs assigned to frameshift substitutions are significantly more conservative than those to random substitutions. The P-values of the t-tests between FSS and RSS are 2.497×10^−10^ (forward frameshifting vs random substitutions) and 2.896×10^−9^ (reverse frameshifting vs random substitutions), respectively.

In the most common scoring matrices, such as BLOSSUM62, PAM250, and GON250, most scores are negative, and the percentage of positive scores is about 35%, *i*.*e*., in random codon substitutions, the percent of positive substitution is about 35%, which is consistent with the observed average random similarity, 0.383 (Table 2b). However, as mentioned above, the frame similarities of real genes are significantly higher than not only the random similarities but also the fame similarities of random CDSs, implying that the shiftability of genes is determined at two different levels, the genetic code, and the coding sequences.

### 3.3 The natural genetic code ranks at the top of all possible codon tables

To further investigate the frameshift optimality of the genetic code, we compared it with two kinds of alternative codon tables:

1. *Random codon tables* are produced by swapping the amino acids assigned to sense codons while keeping all degenerative codons synonymous (*Freeland & Hurst*) [5]. From all possible (20! = 2.43290201×10^18^) random codon tables, 100 independent samples were sampled using a simple random sampling algorithm, each containing one million random codon tables. As shown in Fig 3 and Table 5, when the FSSs were computed using PAM250, BLOSSUM62, and GON250 scoring matrix, the sum FSS of the standard genetic code ranks in the top 13.26%, 1.98%, and 2.94% of all random genetic codes, respectively. For the 100 independent samples, the standard deviations of the means and the ranks of FSSs are all as low as 0.03-0.15%, indicating that the sample size of one million is sufficient.
2. *Compatible codon tables* are produced by permuting the bases in the three different codon positions independently and preserving the AA assignment (*Itzkovitz & Alon*) [7]. For each codon position, there are 4! = 24 possible permutations of the four nucleotides. All 24^3^ = 13,824 “compatible” codon tables were produced, and their FSSs were computed (Supp S3). As shown in Fig 3 and Table 5, the natural genetic code ranks in the top 30.91% of the compatible genetic codes when their FSSs were computed using the PAM250 scoring matrix but ranks in the top 3.48% when using BLOSSUM62 or GON250.

**Table 5.** The usage of codons and their weighted mean FSSs (Gon250)

| *NO* | *Species* | *Weighted mean FSS* |
| --- | --- | --- |
|  | *(Codon Usage)* |  |
| 1 | *H. sapiens* | -9.82 |
| 2 | *M. musculus* | -13.47 |
| 3 | *X. tropicalis* | -12.75 |
| 4 | *D. rerio* | -20.58 |
| 5 | *D. melanogaster* | -19.43 |
| 6 | *C. elegans* | -23.38 |
| 7 | *A. thaliana* | -22.52 |
| 8 | *S. cerevisiae* | -14.08 |
| 9 | *E. coli* | -28.59 |
| 10 | *Equal usage* | -22.27 |

In either case, the ranks of the natural genetic code computed using BLOSSUM62 and GON250 are highly consistent with each other, the standard genetic code ranks in the top 2.0–3.5% of all alternative codon tables in terms of frameshift tolerability. As pointed out by *Itzkovitz* and *Alon* [7], due to the wobble constraint for base pairing in the third position, only two permutations (the identity permutation and the A↔G permutation) are allowed in the third position. Thus, the genetic code has only 24 × 24 × 2 = 1152 distinct alternatives. Of the 1152 unique codes, only a dozen (or a few dozens) are superior to the natural genetic code in terms of frameshift tolerance. Therefore, we conclude that the genetic code is nearly optimal regarding frameshift tolerance.

### 3.4 The shiftability was further optimized at gene-/genome-level

As abovementioned, shiftability is species-dependent (Table 2b). For some real genes, shiftability is exceptionally high (Table S1b), *e*.*g*., *E. coli ydaE* (*δ=*0.571), human glutenin (*δ=*0.660). In other words, shiftability can be adjusted by gene or genome sequences. As shown in Table 6 and Supp S4, the mean FSS weighted by codon usages in *E. coli, A. thaliana*, and *C. elegans* are lower than expected (the mean FSSs of equal usage of codons), showing that frameshift-resistant codons (FTCs) are not overrepresented in these genomes. The weighted mean FSSs are significantly higher than expected in humans, mice, *Xenopus*, and yeast, suggesting that FTCs are overrepresented in these genomes.

**Table 6.** The usage of codon pairs and their weighted mean FSSs (Gon250)

| NO | species | Number of codon pairs |  |  | Weighted mean FSS |  |  |
| --- | --- | --- | --- | --- | --- | --- | --- |
|  |  | Over-represented | Under-represented | Absent | Over-represented | Under-represented | All |
| 1 | *H. sapiens* | 1573 | 2523 | 50 | -1.52 | -7.80 | -3.06 |
| 2 | *M. musculus* | 1505 | 2591 | 190 | -2.83 | -7.13 | -3.81 |
| 3 | *X. tropicalis* | 1660 | 2436 | 148 | -3.12 | -6.98 | -3.80 |
| 4 | *D. rerio* | 1493 | 2603 | 148 | -4.87 | -6.09 | -5.18 |
| 5 | *D. melanogaster* | 1418 | 2678 | 140 | -5.33 | -5.86 | -5.02 |
| 6 | *C. elegans* | 1469 | 2627 | 164 | -6.47 | -5.26 | -6.11 |
| 7 | *A. thaliana* | 1566 | 2530 | 15 | -6.30 | -5.35 | -6.37 |
| 8 | *S. cerevisiae* | 1493 | 2603 | 159 | -4.86 | -6.14 | -4.27 |
| 9 | *E. coli* | 1389 | 2707 | 197 | -6.76 | -5.11 | -6.82 |
| 10 | Equal Usage | 0 | 0 | 0 | N/A | N/A | -5.67 |

On the other hand, frameshifting involves adjacent codon pairs, so the usages of codon pairs are more likely to be related to the shiftability of genes. As shown in Table 7 and Supp S5, the usages of codon pairs are also highly biased in all species tested. Surprisingly, of the 4096 codon pairs, less than 1/3 (⩽1660) are overrepresented, while the remaining (>2400) codon pairs are underrepresented or even unused, suggesting that the synonymous codon pairs had undergone a strong selection pressure [40]. The weighted mean FSSs in *E. coli, C. elegans*, and *A. thaliana* are significantly lower than expected (the mean FSS of equal usage of codon pairs), showing that frameshift-resistant codon pairs (FTCPs) are not overrepresented in these genomes; the weighted mean FSSs are significantly higher than expected in humans, mice, *Xenopus*, and yeast, indicating that FTCPs are overrepresented in these higher species. In these species, shiftability is also higher (Table 2b), suggesting that shiftability is related to the usage of codons and codon pairs.

## 4. Discussion

### 4.1 The shiftability of the genetic code and the coding genes

The natural genetic code has existed since the life origin and is believed to have been optimizing by sense codon reassignment and competition with alternative codes [41]. The natural genetic code was optimized along with several properties during the early history of evolution [42]. It has been reported that the natural genetic code was optimized for the minimization of translational errors, which is explained by the selection to minimize the deleterious effects of translation errors [3]. Besides, it was suggested that only one in every million alternative genetic codes is more efficient than the standard genetic code in terms of minimizing the effects of point-mutations or translational errors [5]; Also, it was shown that the genetic code is nearly optimal for storing additional information within coding sequences, such as out-of-frame hidden stop codons (HSCs) [7].

A complete frameshift is usually a loss of function, and most functional frameshifts are partial frameshifts. Shiftability cannot guarantee that all frameshifts function, but can bring a better chance of restoring normal structure and function in repairing a frameshift mutation [43]. Because of the shiftability, near half of the amino acids remain conserved in a frameshift, regardless of where the frameshifting starts and ends. It is conceivable that a genetic code with greater shiftability had a better chance of winning the competition with its competitors in earlier evolutionary history.

In the above, it is demonstrated that the genetic code guaranteed that, on average, about 40 to 45% of the amino acids are kept conservative in a frameshift. This intriguing property of the genetic code forms the basis of frameshift tolerance, which explains why functional frameshifts could exist [16-18]. If a frameshift is not removed by selecting against, it can be repaired by a reverse mutation, or changed by point mutations [44]. Proteins have been evolving through point and frameshift mutations in their CDSs. The point mutation rate is extremely low so that they alter the sequence, the structure, and the function of proteins at a slow rate. However, frameshift + point mutations provide a far more effective means for a fast-evolving of protein sequences, allowing the emerging of novel or overlapping genes. In the evolutionary process, shiftability can play a vital role in maintaining, repairing, and evolving proteins and coding genes.

### 4.2 The usage of codons and codon pairs

There have been quite some hypotheses on the cause and consequence of the usages of codons/codon pairs, such as gene expression level [45], mRNA structure [46], mRNA stability [47], and protein abundance [48]. Here we demonstrated that the shiftability of a gene or a genome is adjusted through the usage of codons and codon pairs, suggesting that many genes and certain genomes were optimized for frameshift tolerance. The shiftability of coding genes could either be a cause or a consequence of the usage of codons or codon pairs. The more a frameshift resembles the wild type, the more likely it can restore a normal function when a frameshift mutation occurs. Thus, overuse of frameshift-resistant codons or codon pairs confers an evolutionary or survival advantage on a gene or genome. In other words, the frameshift optimality of the genetic code is reflected not only in the code itself but more importantly, in its allowance of further optimizing the frameshift tolerance of a particular gene or genome, sheds light on the role of frameshift mutations in molecular and genomic evolution.

### 4.3 The statistics for measuring frameshift tolerability

We devised a new statistic for frameshift tolerance, frameshift substitution scores, and proved that they are higher in frameshift than in random substitutions. Since we disclosed this study, two other groups have further analyzed the genetic code [27] and proteins [21] using the physicochemical properties (PCPs) of the amino acids. From a chemical point of view, PCP is more suitable for analyzing frameshift tolerance, while FSS could be more convenient in biological studies. Substitution scores are calculated from the probability that different amino acids were substituted by each other over time. Although the substitution scores are ultimately defined by the physicochemical properties of amino acids, their values also reflect the evolutionary relationships of organisms. As such, they are widely used in sequence analyses, *e*.*g*., calculating similarities, constructing alignments, and searching databases.

Each family of scoring matrices has different members, such as PAM1, …, PAM100, and PAM250, representing probabilities of substitution over different timescales. Different scoring matrix members are designed for different evolutionary distances, *e*.*g*., PAM1, …, PAM100 are more suitable for aligning closely related protein sequences, while PAM250 is more suitable for remotely related sequences. Pearson pointed out that “deep” scoring matrices (like BLOSUM62) target alignments with 20 – 30% identity, while “shallow” scoring matrices (*e*.*g*., VTML10), target alignments that share 90 – 50% identity, reflecting much less evolutionary change [49]. The alignment of frameshifts, however, is unique and special, because a frameshift and its wild-type CDS are closely related, their translations have a low identity but a moderate similarity. Obviously, “deep” matrices are more suitable than “shallow” matrices for aligning and analyzing frameshifts. In this study, we adopted three representatives of the “deep” matrices to calculate FSSs. Since frame similarities are quasi-constant, these scoring matrices were used without considering divergence levels. However, it is undetermined which family (or a member of a family) is the most suitable for calculating frameshift tolerance, or whether a specialized scoring matrix specifically designed for analyzing frameshift mutations is needed.

### 4.4 The readthrough rules and their impact on the computation of similarity

In the present study, we incorporate computational frameshifting and readthrough into the analysis. It is important to note that computational frameshifting and readthrough are conceptually different from biological frameshifting and translational readthrough. This does not require that they truly occur in an organism, because these operations are used only for calculating similarities. So, in the present study, they are not taken as biological laws, but computational methods borrowed from biology.

However, in the frameshifts, the expected percentage of stop signals is 3/64 = 4.69%. In real genes, the predicted percentage of hidden stop codons is even higher [8]. Therefore, the readthrough rules can have a significant impact on the frame similarity calculations.

We have conducted a series of studies and found that the location and distribution of hidden stop codons in real genes and their matching wild-type amino acids are not random, and therefore, differences between readthrough and non-readthrough translations are not negligible. All these data suggest that the readthrough rules are probably be adapted to the genetic code and explain part of its optimality. As the presentation of these results depends on the present article, we will present these data in another article.

## 5. Conclusion

Previous studies have proved the robustness of the genetic code to point mutations, and here we analyzed the tolerability of the genetic code and some organisms to frameshift mutations. Based on the above analysis, we conclude that the genetic code and the genomes were both optimized for frameshift tolerance. Shiftability guarantees a near-half similarity of wild types and frameshifts, endowing coding genes an inherent tolerability to frameshift mutations in either (forward or reverse) direction. Thanks to this unique property, the natural genetic code obtained better fitness than its competitors in early evolution. The shiftability serves as an innate mechanism by which genes and genomes tolerate frameshift mutations, and thus, deleterious frameshift mutations could have been utilized as a driving force for evolution. However, the impacts of frameshift tolerance on molecular or genomic evolution remain to be characterized across the tree of life.

## Data accessibility

The source code of the java programs used to analyze the data are available at GitHub (https://github.com/CAUSA/Frameshift). The Supplementary datasets are available at FigShare (https://doi.org/10.6084/m9.figshare.9948050.v2). S1a: Frame similarities aligned by *ClustalW* or *MSA*; S1b: Frame similarities aligned by *FrameAlign*; S2: FSSs of the natural genetic code; S3: FSSs of the alternative genetic codes; S4: FSSs of different codon usages; S5: FSSs of different usages of codon pairs.

## Authors’ Contributions

X. Wang conceived the study, wrote the codes, analyzed the data, prepared the figures and tables, and wrote the paper. Y. Liu and Y. Cai analyzed data. Q. Dong, G. Chen, and J. Zhang discussed the paper and gave suggestions.

## Acknowledgments

This study was funded by the *National Natural Science Foundation of China* through Grant 31571369. We appreciate Dr. Hervé Seligmanna and other anonymous reviewers for their thoughtful comments and suggestions on earlier versions of this paper.

## References

1. Nirenberg, M.W. and J.H. Matthaei, The dependence of cell-free protein synthesis in E. coli upon naturally occurring or synthetic polyribonucleotides. Proc Natl Acad Sci U S A, 1961. 47: p. 4 1588–602.

2. Jukes, T.H. and S. Osawa, Evolutionary changes in the genetic code. Comp Biochem Physiol B, 1993. 106(3): p. 489–94.

3. Haig, D. and L.D. Hurst, A quantitative measure of error minimization in the genetic code. J Mol Evol, 1991. 33(5): p. 412–7.

4. Alff-Steinberger, C., The genetic code and error transmission. Proc Natl Acad Sci U S A, 1969. 64(2): p. 584–91.

5. Freeland, S.J. and L.D. Hurst, The genetic code is one in a million. J Mol Evol, 1998. 47(3): p. 238–48.

6. Guilloux, A. and J.L. Jestin, The genetic code and its optimization for kinetic energy conservation in polypeptide chains. Biosystems, 2012. 109(2): p. 141–4.

7. Itzkovitz, S. and U. Alon, The genetic code is nearly optimal for allowing additional information within protein-coding sequences. Genome Res, 2007. 17(4): p. 405–12.

8. Seligmann, H. and D.D. Pollock, The ambush hypothesis: hidden stop codons prevent off-frame gene reading. DNA Cell Biol, 2004. 23(10): p. 701–5.

9. Loughran, G., et al., Evidence of efficient stop codon readthrough in four mammalian genes. Nucleic Acids Res, 2014. 42(14): p. 8928–38.

10. Jungreis, I., et al., Evidence of abundant stop codon readthrough in Drosophila and other metazoa. Genome Res, 2011. 21(12): p. 2096–113.

11. Schueren, F. and S. Thoms, Functional Translational Readthrough: A Systems Biology Perspective. PLoS Genet, 2016. 12(8): p. e1006196.

12. Chen, J., et al., Dynamic pathways of -1 translational frameshifting. Nature, 2014. 26 512(7514): p. 328–32.

13. Antonov, I., et al., Identification of the nature of reading frame transitions observed in prokaryotic genomes. Nucleic Acids Res, 2013. 41(13): p. 6514–30.

14. Morris, D.K. and V. Lundblad, Programmed translational frameshifting in a gene required for yeast telomere replication. Curr Biol, 1997. 7(12): p. 969–76.

15. Russell, R.D. and A.T. Beckenbach, Recoding of translation in turtle mitochondrial genomes: programmed frameshift mutations and evidence of a modified genetic code. J Mol Evol, 33 2008. 67(6): p. 682–95.

16. Raes, J. and Y. Van de Peer, Functional divergence of proteins through frameshift mutations. Trends Genet, 2005. 21(8): p. 428–31.

17. Pai, H.V., et al., A frameshift mutation and alternate splicing in human brain generate a functional form of the pseudogene cytochrome P4502D7 that demethylates codeine to morphine. J Biol Chem, 2004. 279(26): p. 27383–9.

18. Vandenbussche, M., et al., Structural diversification and neo-functionalization during floral MADS-box gene evolution by C-terminal frameshift mutations. Nucleic Acids Res, 2003. 31(15): p. 4401–9.

19. Hahn, Y. and B. Lee, Identification of nine human-specific frameshift mutations by comparative analysis of the human and the chimpanzee genome sequences. Bioinformatics, 2005. 21 Suppl 1: p. i186–94.

20. Claverie, J.M., Detecting frame shifts by amino acid sequence comparison. J Mol Biol, 1993. 234(4): p. 1140–57.

21. Bartonek, L., D. Braun, and B. Zagrovic, Frameshifting preserves key physicochemical properties of proteins. Proc Natl Acad Sci U S A, 2020. 117(11): p. 5907–5912.

22. Huang, X., et al., Frame-shifted proteins of a given gene retain the same function. Nucleic Acids Res, 2020. 48(8): p. 4396–4404.

23. Diamond, M.E., et al., Overlapping genes in a yeast double-stranded RNA virus. J Virol, 1989. 63(9): p. 3983–90.

24. Chen, N.Y. and H. Paulus, Mechanism of expression of the overlapping genes of Bacillus subtilis aspartokinase II. J Biol Chem, 1988. 263(19): p. 9526–32.

25. Huvet, M. and M.P. Stumpf, Overlapping genes: a window on gene evolvability. BMC Genomics, 2014. 15: p. 721.

26. Wang, X., et al., The shiftability of protein coding genes: the genetic code was optimized for frameshift tolerating. January 23, 2015, PeerJ PrePrints PeerJ.

27. Geyer, R. and A. Madany Mamlouk, On the efficiency of the genetic code after frameshift mutations. PeerJ, 2018. 6: p. e4825.

28. Dabrowski, M., Z. Bukowy-Bieryllo, and E. Zietkiewicz, Translational readthrough potential of natural termination codons in eucaryotes--The impact of RNA sequence. RNA Biol, 2015. 12(9): p. 950–8.

29. Hoffman, E.P. and R.C. Wilhelm, Genetic mapping and dominance of the amber suppressor, Su1 (supD), in Escherichia coli K-12. J Bacteriol, 1970. 103(1): p. 32–6.

30. Kuriki, Y., Temperature-sensitive amber suppression of ompF’-’lacZ fused gene expression in a supE mutant of Escherichia coli K12. FEMS Microbiol Lett, 1993. 107(1): p. 71–6.

31. Johnston, H.M. and J.R. Roth, UGA suppressor that maps within a cluster of ribosomal protein genes. J Bacteriol, 1980. 144(1): p. 300–5.

32. Prather, N.E., B.H. Mims, and E.J. Murgola, supG and supL in Escherichia coli code for mutant lysine tRNAs+. Nucleic Acids Res, 1983. 11(23): p. 8283–6.

33. Chan, T.S. and A. Garen, Amino acid substitutions resulting from suppression of nonsense mutations. V. Tryptophan insertion by the Su9 gene, a suppressor of the UGA nonsense triplet. J Mol Biol, 1970. 49(1): p. 231–4.

34. Henikoff, S. and J.G. Henikoff, Amino acid substitution matrices from protein blocks. Proc Natl Acad Sci U S A, 1992. 89(22): p. 10915–9.

35. Dayhoff, M.O., Computer analysis of protein evolution. Sci Am, 1969. 221(1): p. 86–95.

36. Dayhoff, M.O., The origin and evolution of protein superfamilies. Fed Proc, 1976. 35(10): p. 2132–8.

37. Schneider, A., G.M. Cannarozzi, and G.H. Gonnet, Empirical codon substitution matrix. BMC Bioinformatics, 2005. 6: p. 134.

38. Gutman, G.A. and G.W. Hatfield, Nonrandom utilization of codon pairs in Escherichia coli. Proc Natl Acad Sci U S A, 1989. 86(10): p. 3699–703.

39. Gupta, S.K., J.D. Kececioglu, and A.A. Schaffer, Improving the practical space and time efficiency of the shortest-paths approach to sum-of-pairs multiple sequence alignment. J Comput Biol, 1995. 2(3): p. 459–72.

40. Tats, A., T. Tenson, and M. Remm, Preferred and avoided codon pairs in three domains of life. BMC Genomics, 2008. 9: p. 463.

41. Santos, M.A., et al., Driving change: the evolution of alternative genetic codes. Trends Genet, 2004. 20(2): p. 95–102.

42. Knight, R.D. and L.F. Landweber, The early evolution of the genetic code. Cell, 2000. 101(6): p. 569–72.

43. Wang, X., et al., A frameshift mutation is repaired through nonsense-mediated gene revising in E. coli. bioRxiv, 2020: p. 069971.

44. Dohet, C., R. Wagner, and M. Radman, Methyl-directed repair of frameshift mutations in heteroduplex DNA. Proc Natl Acad Sci U S A, 1986. 83(10): p. 3395–7.

45. Paul, P., A.K. Malakar, and S. Chakraborty, Codon usage and amino acid usage influence genes expression level. Genetica, 2018. 146(1): p. 53–63.

46. Subramanian, A. and R.R. Sarkar, Comparison of codon usage bias across Leishmania and Trypanosomatids to understand mRNA secondary structure, relative protein abundance and pathway functions. Genomics, 2015. 106(4): p. 232–41.

47. Stenoien, H.K. and W. Stephan, Global mRNA stability is not associated with levels of gene expression in Drosophila melanogaster but shows a negative correlation with codon bias. J Mol Evol, 2005. 61(3): p. 306–14.

48. McHardy, A.C., et al., Comparing expression level-dependent features in codon usage with protein abundance: an analysis of ‘predictive proteomics’. Proteomics, 2004. 4(1): p. 46–58.

49. Pearson, W.R., Selecting the Right Similarity-Scoring Matrix. Curr Protoc Bioinformatics, 2013. 43: p. 3.5.1–3.5.9.

